# Inhibition of the Rho/MRTF pathway improves the response of BRAF-resistant melanoma to PD1/PDL1 blockade

**DOI:** 10.1101/2023.12.20.572555

**Authors:** Bardees M. Foda, Sean A. Misek, Kathleen A. Gallo, Richard R. Neubig

**Affiliations:** Department of Pharmacology and Toxicology, Michigan State University, East Lansing, MI, USA; Molecular Genetics and Enzymology Department, National Research Centre, Dokki, Egypt; Department of Physiology, Michigan State University, East Lansing, MI, USA; Broad Institute of MIT and Harvard, Cambridge, MA, 02142, USA; Nicholas V. Perricone, M.D. Division of Dermatology, Department of Medicine, Michigan State University, East Lansing, MI, USA

**Keywords:** Melanoma, Immunotherapy, Resistance, Rho GTPases, MRTF

## Abstract

Metastatic cutaneous melanoma is a fatal skin cancer. Resistance to targeted and immune therapies limits the benefits of current treatments. Identifying and adding anti-resistance agents to current treatment protocols can potentially improve clinical responses. Myocardin-related transcription factor (MRTF) is a transcriptional coactivator whose activity is indirectly regulated by actin and the Rho family of GTPases. We previously demonstrated that development of BRAF inhibitor (BRAFi) resistance frequently activates the Rho/MRTF pathway in human and mouse BRAF^V600E^ melanomas. In clinical trials, pre-treatment with BRAFi reduces the benefit of immune therapies. We aimed to test the efficacy of concurrent treatment with our MRTF pathway inhibitor CCG-257081 and anti-PD1 *in vivo* and to examine its effects on the melanoma immune microenvironment. Because MRTF pathway activation upregulates the expression of immune checkpoint inhibitor genes/proteins, we asked whether CCG-257081 can improve the response to immune checkpoint blockade. CCG-257081 reduced the expression of PDL1 in BRAFi-resistant melanoma cells and decreased surface PDL1 levels on both BRAFi- sensitive and -resistant melanoma cells. Using our recently described murine vemurafenib-resistant melanoma model, we found that CCG-257081, in combination with anti-PD1 immune therapy, reduced tumor growth and increased survival. Moreover, anti-PD1/CCG-257081 co-treatment increased infiltration of CD8^+^ T cells and B cells into the tumor microenvironment and reduced tumor-associated macrophages. Here, we propose CCG-257081 as an anti-resistance and immune therapy-enhancing anti-melanoma agent.

**Novelty and Impact:** We present a study that provides evidence for a new combined approach for targeting BRAF inhibitor-resistant melanoma. Pharmacological inhibition of the resistance-inducing Rho/MRTF pathway using CCG-257081 enhanced the response to PD1/PDL-1 blockade *in vivo*. These results indicate a role of the Rho/MRTF pathway in regulating tumor-immune interactions. Thus, CCG-257081 emerges as a potential new anti-resistance agent that can improve the response to immune checkpoint inhibitors in advanced melanoma and, possibly, other cancers.

## 1 INTRODUCTION

Therapy resistance and immune evasion are common hurdles that limit the efficacy of most anti-cancer therapies. BRAF inhibitors (BRAFi), including vemurafenib (Vem) and dabrafenib, and mitogen-activated protein kinase inhibitors (MEKi), such as trametinib, control melanoma growth in patients with the common BRAF^V600E^ driver oncogene. Most patients respond quickly, but virtually all develop drug resistance.^1–3^ Immune checkpoint therapies (ICTs), including neutralizing antibodies targeting CTLA-4, PDL1, and PD1, block exhaustion signaling of CD8^+^ T cells and prolong T cell effector function.^4–6^ ICTs have greatly improved melanoma outcomes, but they have a slower onset of action compared to targeted therapies.^7–11^ Combined triple therapy using BRAFi, MEKi, and immune checkpoint inhibitors reduced some immune escape mechanisms and improved patient survival.^12,13^ Even with triple combination treatment, however, adverse effects and resistance development prevent optimal outcomes of ICT for many patients.^14,15^

A cross-over comparison of targeted therapy (dabrafenib/trametinib) and ICT (nivolumab/ipilimumab), the DreamSeq study, found that sequencing ICT before targeted therapy improved outcomes.^11^ The reduced effectiveness of ICT after targeted therapy treatment raises the question of whether mechanisms of resistance to targeted therapy also impair ICT responses^16^ and whether blocking resistance to targeted therapy might enhance ICT responses. There are many mechanisms of resistance to BRAF/MEK-targeted therapies.^1,2^ Among them, de-differentiation plays an important role, with contributions by AXL and Rho GTPases along with Rho-driven transcription partners (YAP/TEAD and MRTF/SRF).^17,18^ Here, we explore activation of the Rho/MRTF pathway^18,19^ and its interaction with ICT – anti-PD1 treatment – in a novel isogenic mouse model of targeted therapy resistance.^19^

Rho/MRTF pathway activation occurs in ∼50% of melanomas resistant to targeted therapy.^18^ Cytoskeletal rearrangements and expression of genes involved in proliferation, differentiation, focal adhesion, and migration.^20,21^ There are also interactions between the Rho/MRTF pathway and the immune system. Actin cytoskeleton rearrangement drives immune resistance in breast cancer via concentrating PDL1 at the immunological synapses.^22^ Actin remodeling promotes immune cell-mediated tumor cell killing, including target cell recognition, activation of immune cells, and coordination of immune synapses.^23^ Cell surface rigidity and stiffness affect the interaction between immune and cancer cells in a context-specific manner.^24^

We previously showed that inhibition of the Rho/MRTF pathway re-sensitized BRAFi-resistant melanoma cells to the BRAF inhibitor Vem.^18,19^ Here, we assess the impact of MRTF pathway inhibition on immune responses and tumor progression *in vivo*. The MRTF pathway inhibitor (MRTFi) CCG-257081 reduced transcript and surface protein levels of PDL1 in BRAFi-resistant mouse melanoma cells. These resistant cells also failed to respond to anti-PD1 treatment *in vivo*. Inhibition of the Rho/MRTF pathway augmented the response to anti-PD1 treatment; a combination of CCG-257081 and anti-PD1 antibody reduced tumor growth and increased infiltration of CD8^+^ T-cells and B-cells into the tumor microenvironment. It also decreased infiltration of immunosuppressive M2 macrophages. Overall, this work shows that the MRTF pathway inhibitor, CCG-257081, can enhance the response to anti-PD1 neutralizing antibodies in BRAFi-resistant melanoma *in vivo*.

## 2 MATERIALS AND METHODS

### 2.1 Cell culture

YUMM1.7D4 (RRID: CVCL_ZL99) and YUMMER1.7D4 (RRID: CVCL_A2BD) cell lines were

purchased from Millipore Sigma (YUMM1.7D4: Cat# SCC227; YUMMER1.7D4: Cat# SCC243). These YUMM cells were maintained in the DMEM-F12 medium (ATCC# 30-2006) supplemented with 10% fetal bovine serum (Gibco #10437-028), 1% NEAA (Gibco# 11140-50), and 1% Pen-Strep (ThermoFisher #15140122). YUMM1.7D4 is abbreviated as YUMM1.7 and YUMMER1.7D4 as YUMMER. Vem-resistant YUMM lines, YUMM1.7_R, and YUMMER_R cells were generated as described previously.^19^ All experiments were performed with mycoplasma-free cells. UACC-62 (RRID: CVCL_1780) and SK-MEL-19 (RRID: CVCL_6025) cells were obtained from Dr. Maria Soengas at The University of Michigan; UACC62R cells were made resistant as described previously.^18^ SK-MEL-147 (RRID: CVCL_3876), was previously described.^25^ Cells were routinely tested for mycoplasma contamination by DAPI staining. Short tandem repeat (STR) profiles were performed on SK-Mel-19 and SK-Mel-147 cell lines (Genewiz, South Plainfield, NJ, USA).^26^ Short Tandem Repeat profiling of UACC62 and its isogenic vemurafenib-resistant (UACC62R) cell lines was performed at the MSU genomics core.^18^ All human cell lines have been authenticated using STR profiling within the last three years of generating the RNAseq data.

### 2.2 RNA extraction, cDNA synthesis, and qRT-PCR analysis

Total RNA was isolated using the RNeasy Plus Kit (Qiagen #74134) and was converted to cDNA using the High-Capacity cDNA RT kit (ThermoFisher #4368814) according to the manufacturer’s guidelines. qPCR was performed using the SYBR Green PCR Master Mix (ThermoFisher #4309155) on the Applied Biosystems QuantStudio 7 Flex Real-Time PCR. qPCR primers were designed using the Harvard Primer Bank tool (https://pga.mgh.harvard.edu/primerbank/) and purchased from Integrated DNA Technologies. The primers used in the study included *Tbp* Forward (5’-GCAATGTCTAACGGGGTTTACG -3’), *Tbp* Reverse (5’-TAGAGGTGTGCTGGACACTAC-3’); *Pdl1* Forward (5’- GTCAATGCCCCATACCGCAA-3’); *Pdl1* Reverse (5’-GGCCTGACATATTAGTTCATGCT-3’).

### 2.3 RNA-Seq sample preparation and data processing

Data were collected and processed previously.^18,26^ The results (GEO under the GSE115938) were examined for expression of immune checkpoint-related genes (see details in Supplementary Materials).

### 2.4 Mice and tumor engraftment

Animal studies were performed in accordance with protocols approved by the Michigan State University Institutional Animal Care and Use Committee (IACUC). C57BL/6J mice were purchased from The Jackson Laboratory. Four-week-old male mice were housed at the MSU animal facility for two weeks before initiating animal experiments. Six-week-old mice were shaved and randomized into four groups. YUMMER_R cells were grown to 75% confluence, dissociated with trypsin, washed with sterile PBS (ThermoFisher, 10010023), and resuspended at a density of 1×10^6^ cells in 100 µl PBS. Cells were subcutaneously transplanted into the flank using a 27G needle. Post-injection, mice were monitored for the development of palpable tumors every other day with digital calipers, and tumor volume was computed using the formula (length × width^2^ x 0.5).

### 2.5 Combined treatment of resistant melanoma-bearing mice with anti-PD1 antibody and CCG-257081

Mice with about 200 mm^3^ YUMMER_R tumors were randomized into four groups and treated with (1) isotype control antibody and vehicle, (2) isotype control antibody and CCG-257081, (3) anti-PD1 and vehicle, and (4) anti-PD1 and CCG-257081. For antibody treatment, mice were injected intraperitoneally (ip) every three days with 150 µg of anti-PD1 (clone RMP1-14) or isotype control antibodies (clone 2A3) purchased from BioXcell. Mice received a total of four antibody doses. CCG-257081 was synthesized in the MSU Medicinal Chemistry Core, http://drugdiscovery.msu.edu/facilities/medicinal-chemistry-core-mcc/index.aspx) and delivered daily by ip in a vehicle containing 5% DMSO and 95% polyethylene glycol-400. Tumor size and body weight were recorded every two days; mice were euthanized after 4^th^ dose of anti-PD1 antibody. A humane endpoint, defined as the time for a tumor size of >750 mm^3^, was used for calculating mouse survival.

### 2.6 Flow cytometry analysis of surface PDL1 protein expression on YUMM cells

YUMM cells were seeded in 6-well plates (200K cells per well) and allowed to adhere overnight. Adherent cells were treated with CCG-257081 (3 µM or 10 µM) for 24 hrs. Cells were then dissociated with trypsin, washed in PBS, and suspended in FACS buffer (PBS supplemented with 2% FBS, 0.1% Sodium Azide). To assess cell viability, single-cell suspensions were first stained using the Zombie NIR fixable viability kit (Biolegend, 423105). Next, cells were incubated with FC Block (BD Biosciences, Fc1.3216) and anti-PDL1 (Biolegend, 10F.9G2) in the dark for 30 min at 4°C as the manufacturers recommended. The stained cells were washed twice with FACS buffer, and spectra were analyzed using a 5-laser Cytek Aurora Spectral Cytometer. Spectra were unmixed using SpectroFlo software, and flow cytometric data was analyzed using FlowJo version 10.8.1 (FlowJo, LLC).

### 2.7 Immunophenotyping of tumor microenvironment by flow cytometry

Tumors were isolated and collected in RPMI 1640 supplemented with 2% FBS (Gibco #10437-028) and 1% pen-Strep (ThermoFisher# 15140122). Tumors were digested for 1 hr at 37°C in RPMI medium supplemented with 10% FBS, 1000 units of DNase I-type II (Sigma# D4527), and Collagenase II (Sigma# C1764). Digested tumors were filtered using a 40 μm cell strainer (VWR# 352340), and the digestion reaction was stopped by adding 0.5 μM EDTA. To assess cell viability, single-cell suspensions were first stained using the LIVE/DEAD Fixable Blue Dead Cell Kit (ThermoFisher# L34961) as recommended by the manufacturer. Next, cells were suspended in brilliant buffer (BD Biosciences # 563794) and treated with FC Block (BD Biosciences, Fc1.3216). Cells were stained in the dark for 30 min at 4°C for surface markers using a panel composed of 15 fluorochrome-labeled antibodies specific for CD45 (30-F11), CD4 (RM4– 5), and PD1 (RMP1-30) purchased from ThermoFisher, CD3ε (145–2C11), CD8α (53–6.7), CD19 (ID3), CD11b (M1–70), CD11c (HL3), GR-1 (RB6–8C5), Lag3 (C9B7W), NK1.1 (PK136), F4/80 (BM8), CD206 (C06802), PDL1 (10F.9G2), and Tim3 (RMT3-23) purchased from Biolegend. The stained cells were washed twice with FACS buffer, and spectra were acquired using a 5-laser Cytek Aurora Spectral Cytometer. Spectra were unmixed using SpectroFlo software, and flow cytometric data was analyzed using FlowJo version 10.8.1 (FlowJo, LLC).

### 2.8 Immunohistochemistry

Formalin-fixed paraffin-embedded tumors were sectioned into 4 µm sections, and unstained tissue slides were prepared by the histology core at MSU. The unstained slides were incubated at 55°C overnight. Immunohistochemistry staining was performed following the recommended manufacturer’s protocol for paraffin-embedded tissues. Briefly, tissue sections were deparaffinized, rehydrated, and heated in 10 mM citrate buffer of pH 6.0 to unmask antigens. Inactivation of endogenous peroxidase was carried out by treating the tissues with 3% H_2_O_2_, followed by blocking with 1% BSA and 5% normal goat serum. Tumor sections were incubated overnight with primary antibodies specific for CD8 (Proteintech, 29896-1-AP) or CD206 (Abcam # ab64693). Tumor sections were washed and incubated with goat anti-rabbit IgG, HRP-linked antibody (Cell Signaling, 7074). Subsequent staining procedures were performed following Vector Laboratories’ instructions for the HRP/DAB detection kit (SK-4100). Sections were counterstained with Mayer’s Hematoxylin (Sigma, MH532), then dehydrated and mounted using Permount mounting media (ThermosFisher, SP15-100). Images were acquired using Olympus BX41 Fluorescence Microscope. For quantification, a minimum of five fields of 40x captured images per tissue were analyzed blindly using ImageJ software (version: 2.14.0/1.54f).

### 2.9 Statistical analysis

Statistical analysis was performed as indicated by unpaired two-tailed t-tests for comparative analysis between two groups. A two-way ANOVA test was used for the tumor size analysis. A log-rank test was used for the analysis of mouse survival. Data are presented as mean ± S.E.M, and a p-value < 0.05 was considered statistically significant. All statistical analyses were performed using the GraphPad Prism version 10.1 software (La Jolla, CA).

## 3. RESULTS

### 3.1 Pharmacological inhibition of the MRTF pathway downregulates the expression of PDL1

We previously generated Vemurafenib (Vem)-resistant mouse melanoma cell lines (YUMM1.7_R and YUMMER_R)^19^ and found that they displayed high Rho/MRTF activity and elevated mRNA expression of immune checkpoint inhibitor (ICI) genes, including *Lgals9*, *Ido1, and Pdl1*. We also asked if the expression of ICI genes was increased in human melanoma cells with high Rho/MRTF activities. Consistent with our findings in the resistant YUMM cells, human Vem-resistant UACC62R and the NRAS-mutant SK-Mel-147 melanoma lines, which had high Rho/MRTF signaling, showed upregulation of many ICIs (Supplementary Table 1). Additionally, UACC62R and SK-Mel-147 human lines displayed significant downregulation of genes encoding co-stimulatory immune molecules such as *CD40*, *ICOS*, and *TNFRSF14* compared to their drug-sensitive counterparts (UACC62P and SK-Mel-19 – see Supplementary Table 2).

Given the association of high Rho/MRTF activity with elevated ICI expression, we asked whether pharmacological inhibition of Rho/MRTF could impact the *Pdl1* mRNA or protein expression in our Vem-resistant mouse melanoma cells. We treated parental (YUMMER_P) and resistant (YUMMER_R) cells with 3 µM or 10 µM of CCG-257081. The basal mRNA levels of *Pdl1* in DMSO-treated YUMMER_R cells were 53% higher than in YUMMER_P cells (Figure 1A). CCG-257081 treatment reduced *Pdl1* mRNA levels in YUMMER_R cells in a concentration-dependent manner (Figure 1B) but did not significantly affect *Pdl1* mRNA levels of YUMMER_P cells. These data suggest that this ICI transcriptional upregulation depends at least in part on Rho/MRTF upregulation. To assess if the differences in *Pdl1* mRNA levels in parental and resistant cells are also reflected in cell surface protein expression, flow cytometry was performed. The surface expression of PDL1 protein on YUMMER_P and YUMMER_R cells showed similar median fluorescence intensity (MFI) (Figure 1C and 1D). Interestingly, treatment with 3 µM or 10 µM of CCG-257081 reduced the surface PDL1 protein expression on both YUMMER_P and YUMMER_R cells (Figure 1E-1H). Our results align with a previous report that MRTF-A facilitates gene expression of *Pdl1* in lung cancer cells.^27^ Taken together, these data indicate that inhibition of Rho/MRTF in melanoma cells regulates mRNA and cell surface levels of immune checkpoint proteins, suggesting that the Rho/MRTF pathway may modulate tumor-immune interactions.

**Figure 1:**
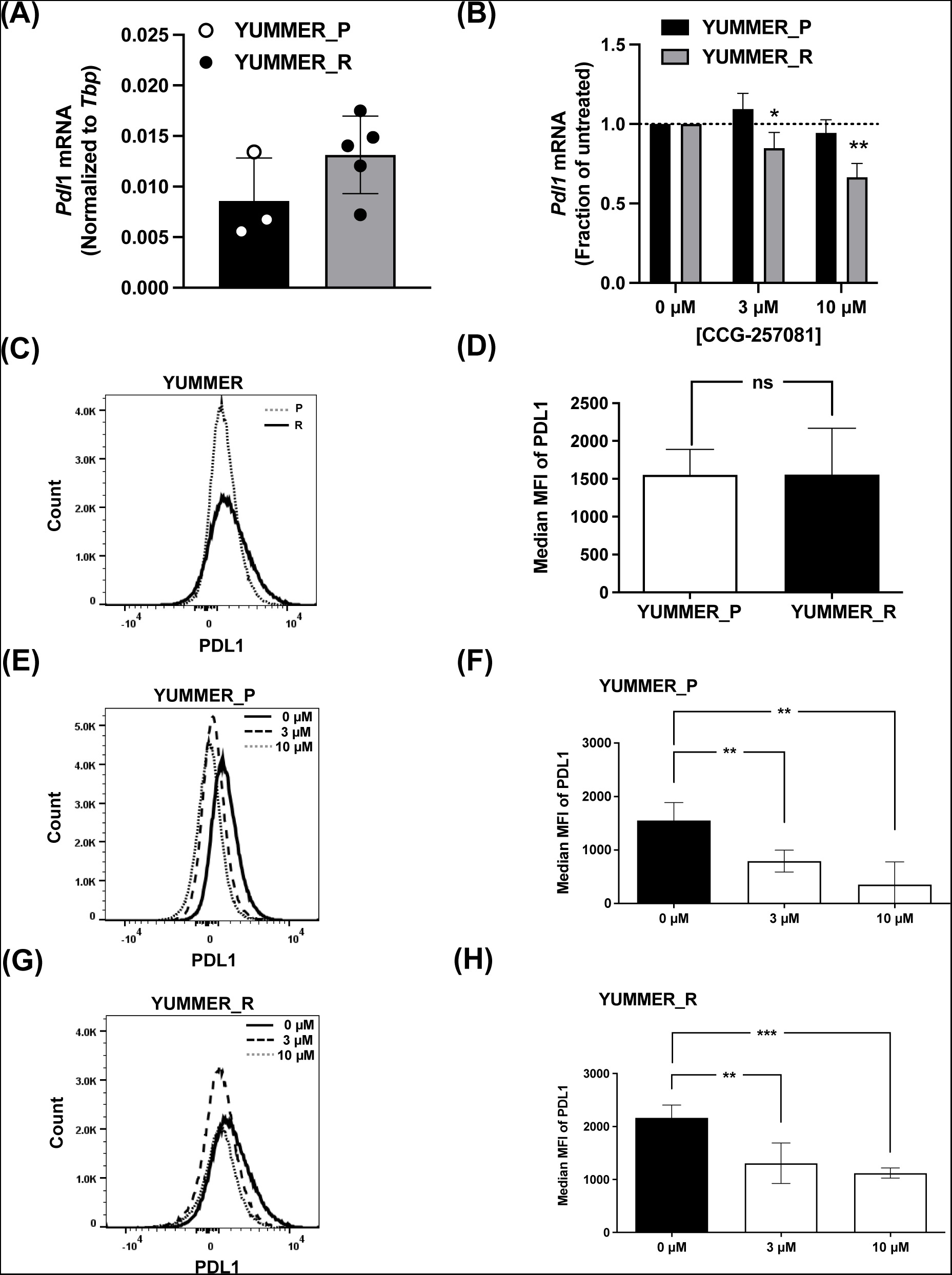
Inhibition of the Rho/MRTF pathway reduces the mRNA expression of *Pdl1* and suppresses surface protein levels on resistant melanoma cells *in vitro*. **(A)** qRT-PCR analysis of *Pdl11* mRNA of YUMMER_P and YUMMER_R cells. Values were normalized to a *Tbp* reference gene. (**B**) YUMMER_P and YUMMER_R cells were treated with the indicated concentrations of CCG-257081 for 24 hr. *Pdl1* mRNA levels are expressed relative to untreated cells. **(C)** Representative flow cytometric profiles of cell surface PDL1 protein of untreated YUMMER_P and YUMMER_R. **(D)** Median fluorescence intensity (MFI) of surface PDL1 for untreated YUMMER cells. Representative flow cytometric profiles of cell surface PDL1 protein of **(E)** YUMMER_P and **(G)** YUMMER_R treated with the indicated concentration of CCG-257081 for 24 hr. (**F, H)** MFI of surface PDL1 of CCG-25708-treated YUMMER_P (F) and YUMMER_R (H). Results are the mean of at least three independent experiments, * *p* < 0.05; ** *p* < 0.01; **** *p* < 0.0001 by unpaired t-test; ns: not significant.

### 3.2 Rho/MRTF inhibition improves the *in vivo* response to the ICT antibody anti-PD1

We utilized YUMMER_R cells *in vivo* to test the impact of concurrent treatment with an anti-PD1 neutralizing antibody and CCG-257081. We intraperitoneally injected YUMMER_R tumor-bearing mice with an anti-PD1 neutralizing antibody alone, CCG-257081 alone, or a combination of the two (Figure 2A). The mice received a minimal dose of anti-PD1 (150 µg every three days) to avoid adverse effects, and they received 100 mg/kg of CCG-257081 in a vehicle containing 95% ethylene glycol 400 and 5% DMSO. YUMMER_R tumors were resistant to single-agent anti-PD1 treatment, as evidenced by no significant reduction in tumor size or change in mouse survival with PD1 blockade (Figure 2B). Similarly, mice treated with CCG-257081 alone showed no change in tumor size (Fig 2B). The combined anti-PD1/CCG-257081 treatment significantly reduced the growth of the YUMMER_R tumor and increased mouse survival compared to no treatment and to each individual treatment (Figure 2C). Additionally, co-treatment reduced skin ulcer formation at the tumor site (Figure 2D and 2E). These results show that pharmacological inhibition of the activated Rho/MRTF pathway can enhance the response of BRAFi-resistant melanoma to ICT in preclinical models.

**Figure 2:**
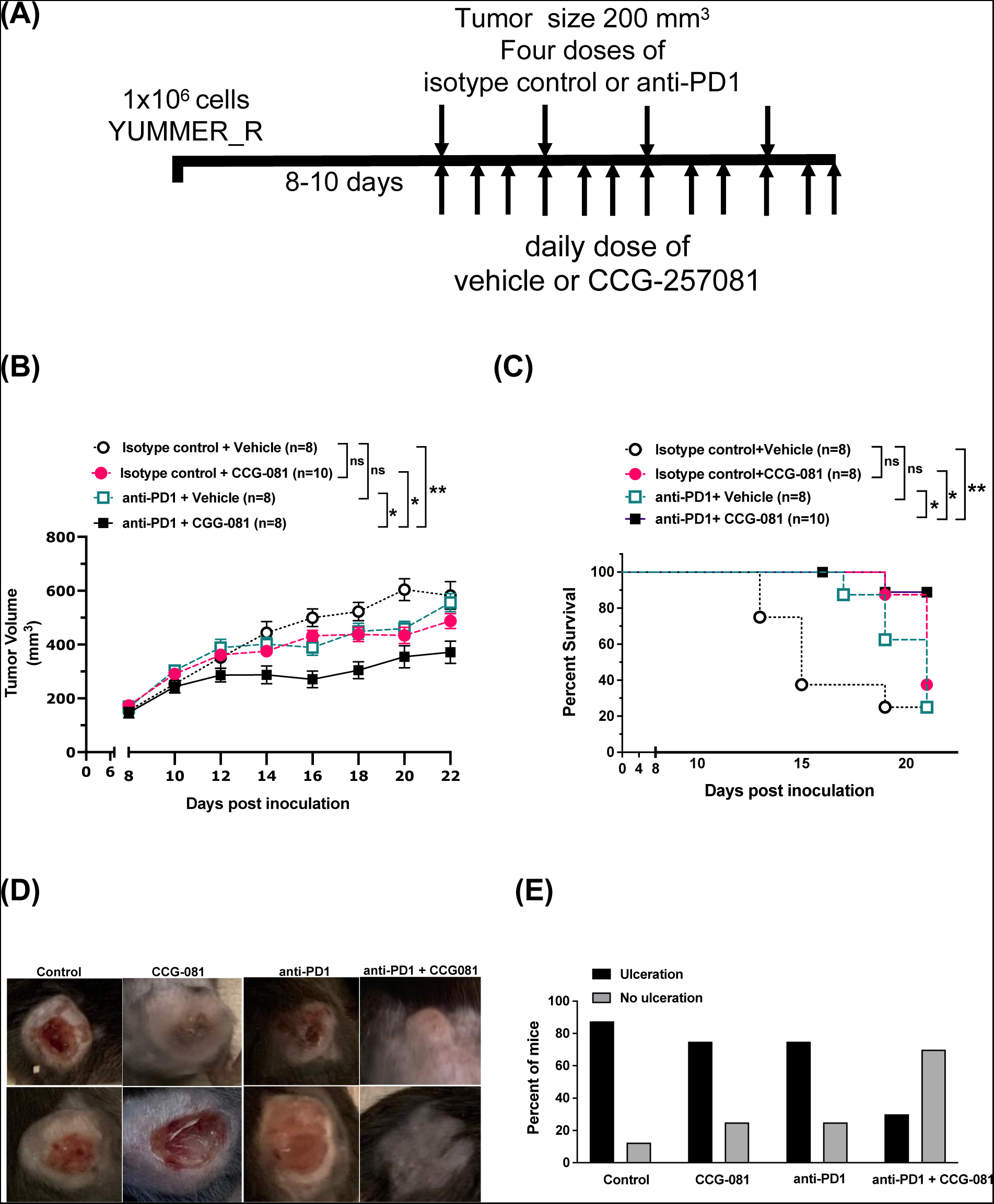
Co-treatment of the YUMMER_R tumor-bearing mice with anti-PD1 and CCG-257081 suppresses tumor growth and extends survival. **(A)** Schematic diagram of the experimental timeline of YUMMER_R tumor development in C57BL/6 mice and treatment strategy. After tumors reached 200 mm^3^, mice were injected intraperitoneally with either 150 µg of anti-PD1 or isotype antibody every three days for 12 days (total of four doses). They also received daily doses of either 100 mg/kg of CCG-257081 or the equivalent volume of vehicle (∼12 doses), resulting in 4 treatment groups. (**B)** Tumor growth was monitored, and a 2-way ANOVA was used for estimating significant differences in tumor size among groups. Results are expressed as the mean ± SEM for the indicated mouse numbers (n); * p < 0.05; ** p < 0.01; ns: not significant. (**C**) Percentage of surviving mice. For the survival curve, the endpoint was defined as the time for a tumor size to reach >750 mm^3^; a log-rank test was used to determine significant differences in survival curves. Results are expressed as the medians for the indicated mouse numbers (n); * p < 0.05; ** p < 0.01; ns: not significant. (**D)** Images of YUMMER_R tumor/skin ulceration from 2 representative mice from each treatment group after the full course of treatment. **(E)** Percentage of mice with and without skin ulceration. CCG-257081 is abbreviated as CCG-081

### 3.3 CCG-257081/anti-PD1 co-treatment increased tumor-infiltrating CD8^+^ T cells and B cells

Across cancer types, CD8^+^ T-cell infiltration is associated with improved responses to immune checkpoint inhibitors.^28^ To better understand the effects of Rho/MRTF pathway inhibition alone and with anti-PD1 treatment on the melanoma immune microenvironment, we used flow cytometry (FC) and immunohistochemistry (IHC) to analyze the immune cell complement in the tumor. A panel of 17 fluorochrome-labeled antibodies was used to identify/interrogate different subtypes of immune cells following an immune gating scheme (Figure S1). In the FC analysis, anti-PD1 alone and in combination with CCG-257081 led to significant increases of CD8^+^ T cells in the tumors (Figure 4B). In the IHC analysis, only mice co-treated with CCG-257081 and anti-PD1 combined showed a statistically significant increase in CD8^+^ T cells (Figure 3A and 3B). In addition to the frequency of cells, the functional state (e.g., exhaustion) of CD8^+^ T cells can be relevant. However, we did not observe any change in the composite exhaustion status of CD8^+^ T cells as assessed by the fraction of PD1^+^ cells that are also positive for Lag3 and Tim1 (Figure S2).

**Figure 3.**
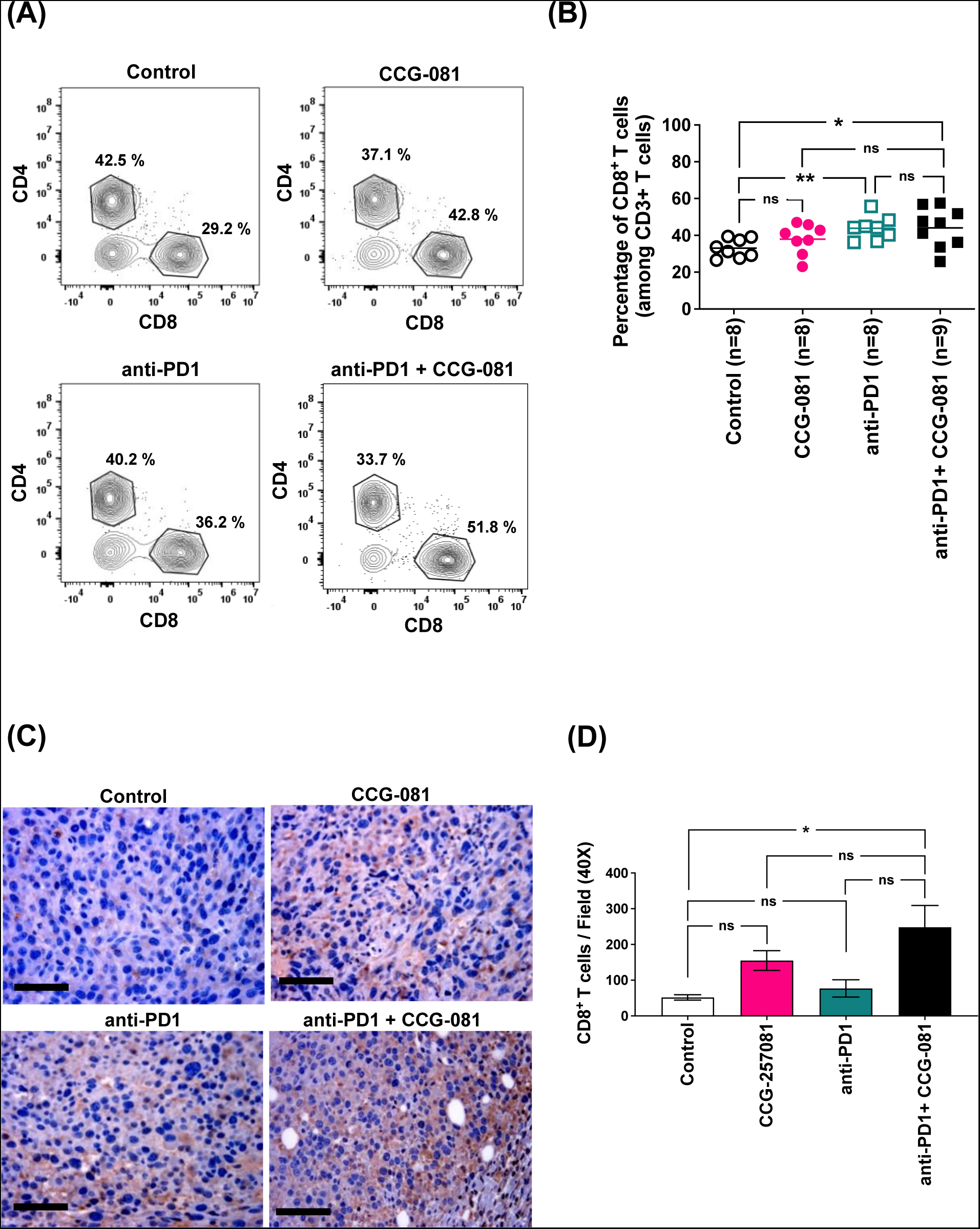
Increased infiltration of CD8^+^ T cells into the tumor microenvironment of the mice co-treated with anti-PD1 and CCG-257081. **(A):** Representative flow cytometry profiles (gated on CD3^+^ T cells). Single-cell suspensions were prepared from tumors isolated from the indicated four mice groups. Cells were analyzed for the frequency of CD8^+^ T cells; single cell suspension was stained with an antibody panel, including antibodies specific for CD45, CD3, CD8, and CD4 and zombie dye for cell viability. **(B):** Percentage of CD8^+^ T cells from the indicated numbers of mice; each data point represents a single mouse; \**p < 0.05* by Mann-Whitney U test; ns: not significant. **(C):** Immunohistochemistry staining of tumor tissues stained with CD8 antibody; scale bar is 5 mm. **(D):** Number of CD8^+^ T cells detected in five fields of tumor from each mouse; \**p < 0.05* by unpaired t-test. CCG-257081 is abbreviated as CCG-081

**Figure 4.**
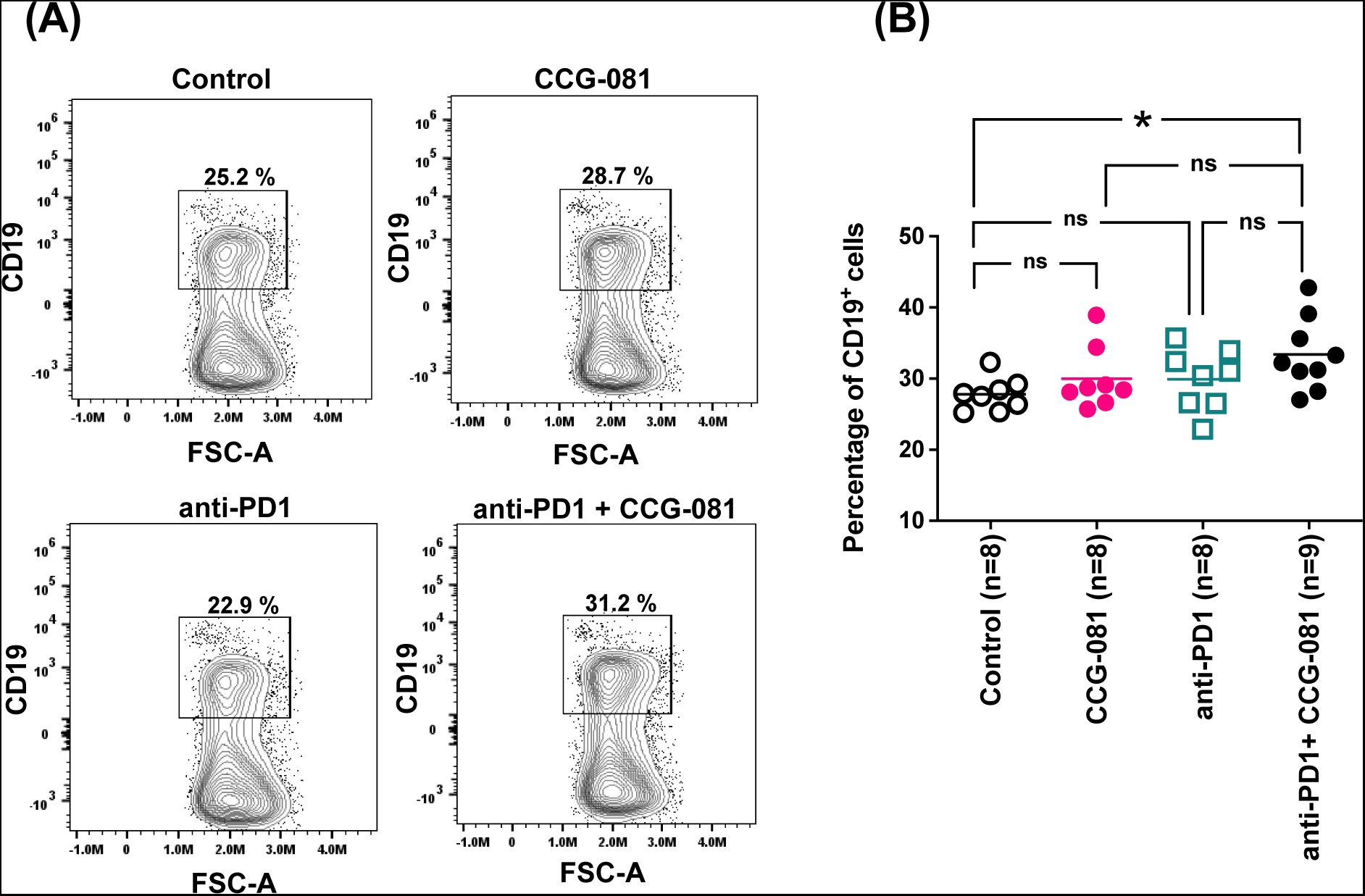
Increased infiltration of B cells into the tumor microenvironment of mice co-treated with anti-PD1 and CCG-257081. **(A):** Representative flow cytometry profiles (gated on CD3^−^ T cells). Single-cell suspensions were prepared from tumors isolated from the indicated four mice groups. Cells were analyzed for the frequency of B cells (CD19^+^); single cell suspension was stained with an antibody panel, including antibodies specific for CD45, CD3, and CD19 and zombie dye for viable cells. **(B):** Percentages of B cells summarized from the indicated numbers of mice, each data point represents one mouse; *\*p < 0.05* by Mann-Whitney U test; ns: not significant. CCG-257081 is abbreviated as CCG-081

Humoral immunity also contributes to immune responses toward immunogenic tumors^29,30^, so we assessed B cell infiltration in the tumors of treated mice (Figure 5A and 5B). Based on flow cytometry, B-cell infiltration was unaffected by single-agent treatment with either CCG-257081 or the anti-PD1 neutralizing antibody, but a significant increase in infiltration of B-cells was observed in the mice co-treated with anti-PD1 and CCG-257081. These results indicate that coupling PD1/PDL1 blockade with Rho/MRTF pathway inhibition may enhance anti-melanoma immunity and modulate the tumor microenvironment into a more inflammatory milieu.

**Figure 5.**
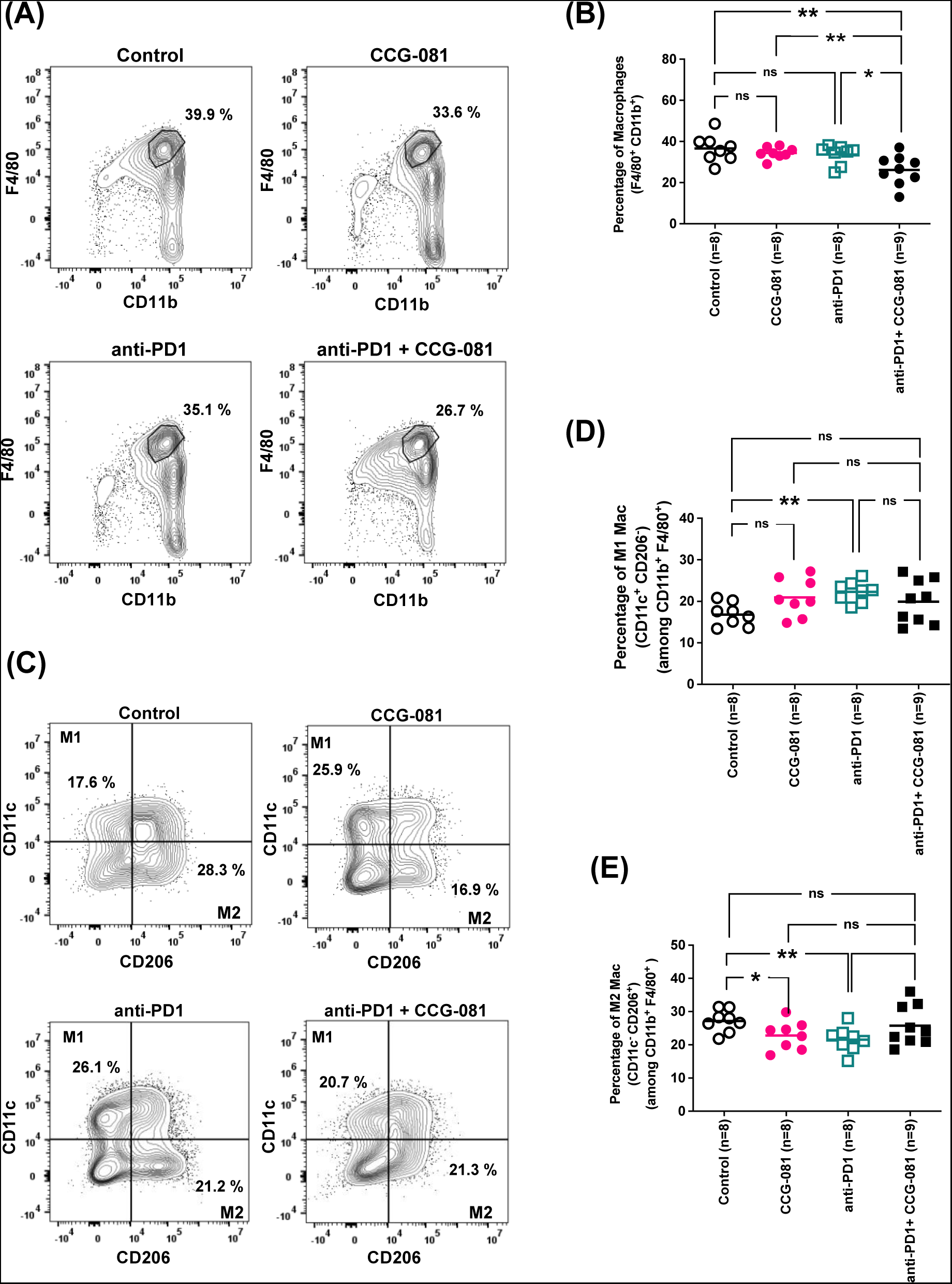
Decreased infiltration of total macrophages and immunosuppressive M2 macrophages into the tumor microenvironment of mice co-treated with anti-PD1 and CCG-257081. **(A):** Representative flow cytometry profiles (gated on CD3^−^ cells). Single-cell suspensions were prepared from tumors collected from the indicated four treated mice groups. Cells were analyzed for the frequency of total macrophages (F4/80^+^ CD11b^+^); single cell suspensions were stained with an antibody panel, including antibodies specific for CD45, CD3, F4/80, and CD11b, as well as zombie dye for viable cells. **(B):** Percentages of total macrophages summarized from the indicated numbers of mice. **(C):** Representative flow cytometry profiles (gated on CD3^−^ F4/80^+^ CD11b^+^ cells). Single-cell suspensions were prepared from tumors collected from the indicated four mouse treatment groups. Tumor cells were analyzed for the frequency of M1 (CD11c^+^ CD206^−^) and M2 macrophages (CD11c^−^ CD206^+^). The single-cell suspension was stained with an antibody panel, including antibodies specific for CD45, CD3, F4/80, CD11b, CD11c, and CD206, as well as zombie dye for viable cells. **(D-E):** Percentages of M1 and M2 macrophages summarized from the indicated numbers of mice; each symbol represents one mouse; *\*p < 0.05; **p < 0.01* by Mann-Whitney U test; ns: not significant. CCG-257081 is abbreviated as CCG-081

**Figure 6.**
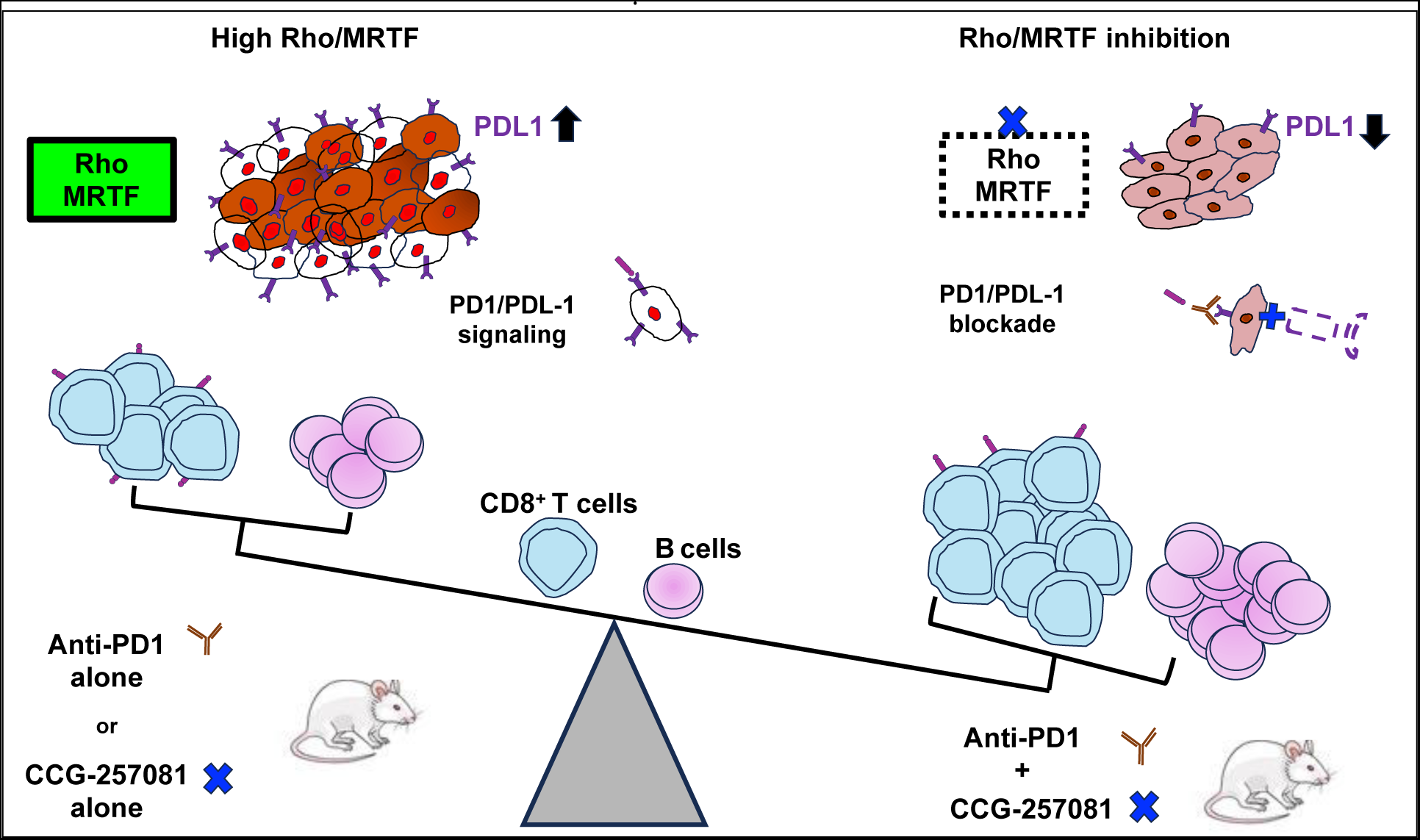
Model of the impact of concurrent inhibition of Rho/MRTF and PD1/PDL1 blockade. Upregulation of the Rho/MRTF pathway in resistant melanoma cells is associated with increased PDL1 expression, aggressive tumor growth, and poor infiltration of CD8^+^ and B cells. Inhibition of the Rho/MRTF pathway by CCG-257081 disrupts the expression of PDL1, allowing anti-PD1 immunotherapy to more effectively block anti-PD1/PDL1 inhibitory signaling. The combination treatment, which improves the response to anti-PD-1 immunotherapy, is associated with an increase in infiltration of CD8^+^ T lymphocytes and B cells into the tumor microenvironment and a reduction in tumor growth. A poorly responded tumor is shown on the left, depicted in red, indicating an aggressive tumor with an activated Rho/MRTF pathway and upregulated PD1/PDL1 signaling. A responsive tumor on the right is displayed in rose color, referring to a less ulcerated tumor with pharmacological blockade of the dysregulated Rho/MRTF and PD1/PDL1.

### 3.4 Rho/MRTF pathway inhibition in conjunction with PD1/PDL1 blockade reduces macrophage tumor infiltration and modulates the macrophage subtype distribution in the melanoma microenvironment

We asked how our combined treatment protocol would affect tumor-associated macrophages (TAMs) in the resistant melanoma tumors by using flow cytometry to assess macrophages (F4/80^+^ CD11b^+^) and macrophage subtypes, M1 (CD11c^+^ CD206^−^) and M2 (CD11c^−^ CD206^+^). While none of the individual treatments reduced the total number of infiltrated macrophages, combined treatment with anti-PD1 and CCG-257081 produced a significant reduction in total TAMs compared to all the other groups (Figure 5A and 5B). We further analyzed the macrophage subtypes inflammatory M1 and immunosuppressive M2. Interestingly, the treatments showed opposite profiles; there was an increase in M1 and a reduction in M2 type macrophages. Each treatment resulted in a greater fraction of inflammatory M1; however, the change was only statistically significant for the tumors treated with anti-PD1 alone (Figure 5C and 5D). The fraction of immunosuppressive M2 (Figure 5C and 5E) decreased significantly with the single treatment of CCG-257081 or anti-PD1 treatment. The combined treatment did not reduce the fraction of M2 macrophages; however, with the decrease in total macrophages, the absolute number of M2 macrophages would be reduced. These results indicate that inhibition of Rho/MRTF, along with an immune checkpoint treatment, can regulate the innate inflammatory milieu within melanoma tumors.

## 4 DISCUSSION

In melanoma treatment, targeted therapies such as BRAFi and MEKi are effective in patients with tumors carrying the prevalent BRAF^V600^ mutations, but responses are not durable due to rapid development of clinical resistance. ICT evokes powerful responses that can be very durable, but not all patients respond. In the DreamSeq study^11^ with combined nivolumab (anti-PD-1)/ipililamab (anti-CTLA4) treatment, the overall survival was only 65% at 5 years, and progression-free survival was only 30-40%. Also, these combination ICT treatments have significant side effects.^11^ The fact that there may be shared resistance mechanisms for targeted drugs and ICT (e.g., increased activation of Rho and its downstream transcription signals) suggests a potential approach to improving melanoma treatment.

Here, we present *in vitro* results and a preclinical study exploring the effects of Rho/MRTF pathway inhibition on ICI gene and protein expression and enhancement of ICT treatment responses. The lack of an effective ICT response in the BRAFi-resistant mouse melanoma lines provides an intriguing correlate to the DreamSeq study where BRAFi/MEKi-pretreatment, and presumably development of resistance, rendered ICT less effective. Similar results have been found in pre-clinical studies in other MAPK-pathway-driven cancers where efficacy is greater when ICT is given prior to targeted therapy.^31^ The ability of CCG-257081, the Rho/MRTF pathway inhibitor, to enhance the ICT response (the current study) as well as to prevent the onset of Vem resistance^19^, to reduce metastasis^25^, and to partially restore Vem sensitivity to resistant melanoma cells^18,19^, suggests broad actions of this compound in improving melanoma treatment.

Highly immunogenic melanoma cells can recruit molecular mechanisms to suppress immunosurveillance.^32,33^ In this study, we found that the Rho/MRTF pathway inhibitor, CCG-257081, could disrupt surface PDL1 protein expression (Figure 1). This is consistent with the observation that the Rho-associated protein kinase (ROCK) stabilizes PDL1 protein levels in breast cancer.^34^ Additionally, actin cytoskeleton rearrangement substantially drives immune resistance in breast cancer via concentrating PDL1 at the immunological synapsis.^22^ We have previously shown that CCG-257081 disrupts actin stress fiber rearrangements in resistant mouse melanoma YUMM cells^19^ These effects of CCG-257081 may, in part, relieve the inhibitory signals transmitted from Vem-resistant tumor cells to immune cells. Further analysis of the involvement of the Rho/MRTF pathway in melanoma-immune interactions would provide deeper insights.

In melanoma, tumor-infiltrating B cells are necessary for preserving inflammation and enhancing immune responses to immune checkpoint blockade.^35,36^ The efficacy of the combined MRTFi/ICT treatment demonstrated here is accompanied by enhanced B cell and CD8^+^ T cell content in the tumors. The observed effects may occur through MRTFi-mediated suppression of melanoma cell ICI expression or through direct actions on immune cells. Whether Rho/MRTF modulates the melanoma immune response by acting directly on melanoma cells and/or on cells in the immune microenvironment warrants future study. In addition, deciphering the specific role of Rho/MRTF in promoting the activation and recruitment of T cells and the maturation and inflammatory functions of B cell subsets deserves examination.

Polarization of M2-TAMs contributes to driving melanoma growth and is associated with metastatic human melanoma.^37,38^ There is a growing interest in macrophage-based therapies to target the functional plasticity of macrophages via depletion or repolarization of M2-TAMs.^39^ There is some prior information on the role of MRTF in macrophage actions. ROCK-mediated cytoskeletal rearrangements increase immunosuppressive M2-TAMs^40^ and MRTF-A modulates the proliferation and apoptosis of macrophages^41^. This could contribute to the decreased levels of total macrophages in the tumor microenvironment upon treatment with CCG-257081. Why only combination therapy with MRTFi and ICT, and not monotherapy, reduced tumor macrophage content is not clear. The combined treatment with MRTFi and ICT increased M1-type macrophages more than any individual treatment, but the effects on M2 macrophage differentiation are less clear. While the percentage of tumor-associated M2 macrophages in the combination treatment was not lower than the others, it is possible that the combination of reduced total macrophages and a similar M2 percentage could contribute to the enhanced ICT response. In line with this, we previously reported that inhibition of the Rho/MRTF pathway in pancreatic cancer reduced the infiltration of macrophages.^42^Hence, more detailed information on the mechanism of the regulatory role of CCG-257081 on macrophage differentiation and recruitment would be of interest to optimize effects in different types of cancers and other auto-immune diseases.

The availability of highly effective anti-resistance melanoma therapies is critical, especially for the subset of patients who do not respond to current immunotherapies. We have previously shown that the Rho/MRTF pathway inhibitor CCG-257081 and related analogs can markedly reduce melanoma metastasis^25^ as well as reverse and even prevent resistance to targeted therapies like BRAFi and MEKi.^19,25,26^ The results presented here suggest that CCG-257081 can also enhance immunotherapy responses. This represents an important opportunity to use a single compound to attack resistance to two of the most important classes of melanoma therapies. Further studies with triple combinations of ICT, targeted therapies, and blockade of the Rho/MRTF pathway may provide a valuable addition to melanoma therapeutics.

## AUTHOR CONTRIBUTIONS

**Bardees Foda** contributed to Conceptualization, Methodology, Formal analysis, Investigation, Writing—Original Draft, and Writing—Review and Editing. **Sean Misek** contributed to Methodology, Review, and Editing. **Kathleen Gallo** contributed to Conceptualization, Review, and Editing. **Richard Neubig** contributed to Conceptualization, Formal analysis, Writing— Review and Editing, Supervision, and Funding acquisition. All authors have read and agreed to the published version of the manuscript. The work reported in the paper has been performed by the authors unless clearly specified in the text.

## ACKNOWLEDGMENT

We thank Drs. Matthew Bernard, Yasser Aldhman, and Maja Blake for their suggestions on the panel design for the Aurora spectral cytometer. We acknowledge Dr. Karen Liby and Dr. Ana Sofia Mendes Leal for their recommendations on IHC. We thank Dr. Daniel Isaac for the helpful discussions on current melanoma treatments. We thank Jeffrey Leipprandt for assistance with the mouse colony and IACUC protocols.

## DTATA AVAILABILITY STATEMENT

The RNA-seq data included in the article is available through the GEO: GSE115938 and GSE134320 repository.^18^ Other data supporting our study’s findings are available from the corresponding author upon request.

## ETHICS STATEMENT

Animal studies were performed in accordance with the International Animal Care and Use Committee (IACUC)-protocols approved by the Michigan State University.

## Abbreviations

BRAFi: BRAF inhibitor
CF: Flow Cytometry
ICIs: Immune checkpoint inhibitor proteins
ICT: Immune checkpoint therapy
IHC: Immunohistochemistry
MAPK: Mitogen-activated protein kinase
MEKi: MAPK inhibitors
MFI: Median fluorescence intensity
MRTF: Myocardin related transcription factor
MRTFi: MRTF pathway inhibitor
PD1: Programmed Cell death protein 1
PDL1: Programmed Cell death Ligand 1
ROCK: Rho-associated protein kinase
TAMs: tumor-associated macrophages
Vem: Vemurafenib
YUMM: Yale University Mouse Melanoma
YUMMER: YUMM exposed to radiation
YUMMER_P: Parental YUMMER (Vem-Sensitive)
YUMMER_R: Resistant YUMM (Vem-Resistant)

