## supplementary data for "Inhibition of the Rho/MRTF pathway improves the response of BRAF-resistant melanoma to PD1/PDL1 blockade"

### 1 SUPPLEMENTARY MATERIALS

#### 1.2 RESULTS

##### 1.2.1 BRAFi and NRAS human melanoma cells are programmed to regulate the expression of several immune checkpoint genes

We previously performed RNSseq analysis for two human melanoma cell lines UACC62R, and SK-Mel-19, which show high Rho/MRTF activity.[1,2] Cellular and molecular analysis show resistance to targeted drug treatment *in vitro* (BRAFi and MEKi, respectively). They also had increased actin stress fibers and upregulated Rho/MRTF-dependent gene expression. Here, we revisited the RNAseq results (GEO: GSE115938 and GSE134320) to assess the expression of immune checkpoint genes relative to their drug-sensitive counterpart cell lines, which displayed lower Rho/MRTF activity (Supplementary Table 1 and Table 2). The human melanoma cells with highly activated Rho/MRTF showed upregulation of many immune checkpoint inhibitor genes; they also showed downregulated immune markers for costimulatory checkpoints.

##### 1.2.2 Effect of combined treatment on mouse exhaustion markers *in vivo*.

Continued immune exposure to cancer cells like YUMMER cells can lead to $CD8^{+}$ T cell exhaustion due to chronic inflammation.[3] Exhausted $CD8^{+}$ T cells have minimal effector function and can ultimately undergo clonal elimination of the tumor cell-directed immune cells.[4] We measured the expression of exhaustion markers such as PD1, LAG3, and TIM3 on $CD8^{+}$ T cells infiltrating the murine tumor microenvironment (Supplementary Figure S1). Anti-PD1 treatment alone resulted in significantly reduced PD1 surface protein expression by flow cytometry (Supplementary Figure S1A). This is consistent with reports that PD1/PDL1 blockade can reverse T cell exhaustion.[5] However, the CCG-257081 single treatment and anti-PD1/CCG-257081 combined treatment showed similar expression of PD1 on $CD8^{+}$ T cells. Expression of the immune checkpoint inhibitor TIM3 on tumor-infiltrating $CD8^{+}$ T cells increased with CCG-257081 mono-treatment and co-treatment (Supplementary Figure S1B). Analysis of LAG3 expression was similar among all treated tumors (Supplementary Figure S1C). Thus, while CCG-257081 could beneficially modulate immune cell infiltration or state-change in the anti-PD1 treated mice, the individual exhaustion markers do not provide evidence of reduced exhaustion with CCG-257081 treatment.

A more robust assessment of the overall exhaustion status of $CD8^+$ T cells is provided by combining the expression of multiple ICIs.[6] We assessed the magnitude of inhibitory signals on tumor-infiltrating $CD8^+$ T cells by measuring the percentage of $CD8^+$ T cells that were positive for ICIs (TIM3 and LAG3) among $PD1^+$ $CD8^+$ T cells (Supplementary Figure S1D). $CD8^+$ T cells infiltrating tumors that received combined treatment had a similar exhaustion score; however, anti-PD1 mono-treatment showed $CD8^+$ T cells with a higher exhaustion score (p-value= 0.053). Interestingly, the exhaustion score of $CD8^+$ T cells of CCG-257081-treated mice was significantly less than that of the anti-PD1 treated group (p-value= 0.03). These results suggest that blocking PD1/PDL1 extended the activation cycles of $CD8^+$ T cells, eventually leading to exhaustion, while inhibition of the Rho/MRTF pathway may delay T cell exhaustion.

**SUPPLEMENTARY FIGURE LEGENDS:**

**Supplementary Figure S1: Expression profile of immune checkpoint inhibitors of CD8+ T cells infiltrated into YUMMER_R tumor microenvironment. (A-C):** Median fluorescence intensity (MFI) of PD1 (A), Lag3 (B), and Tim3 (C). Cells were isolated from tumors collected from the depicted four mice groups. Expression of surface immune checkpoint inhibitors on $CD8^+$ Tumor T cells was measured by calculating the MFI; single cell suspension was stained with an antibody panel, including antibodies specific for CD45, CD3, CD4, CD8, PD1, Lag3, and Tim3, as well as zombie dye for cell viability. **(D):** Exhaustion status of infiltrated $CD8^+$ T cells. Exhaustion is represented as the percentage of CD8 cells that are positive for three exhaustion markers: PD1, Lag3, and Tim3. $^*p < 0.05$; $^{***}p < 0.001$; by Mann-Whitney U test; ns: not significant. Data are the mean of the indicated numbers of mice; each symbol represents one mouse.

**Supplementary Figure S2: Gating strategy of flow cytometry analysis.** Flow cytometry profiles represent the gating strategy of immune cells.

**Supplementary Table 1: Expression of inhibitory immune checkpoints genes in Rho$^{high}$ human melanoma cells.** RNAseq data of twenty-seven immunosuppressive genes of UACC62R relative to UACC62P and SK-Mel-147 relative to SK-Mel-19

| | **Gene** | **UACC62R** | **SK-Mel-147** |
|---|---|---|---|
| | | (Log$_2$ fold change) | |
| *Mostly upregulated genes* | *CD274 (PDL1)* | -0.5* | 3.1 |
| | *PDCD1LG2 (PDL2)* | 6.2 | 10.0 |
| | *IDO1* | 7.7 | 4.9 |
| | *LGALS9* | 6.0 | 1.8 |
| | *TNFRSF9* | 6.1 | 8.5 |
| | *TDO2* | 3.6 | 3.2 |
| | *VSIR* | 2.8 | -3.7* |
| | *NRP1* | 2.9 | 8.9 |
| | *CD96* | 2.1 | 3.8 |
| | *CEACAM1* | -0.4 | 10.0 |
| | *HHLA2* | 0.6* | 0.0* |
| | *CD47* | 0.2* | 0.5 |
| | *CD209* | 0.0* | 0.9 |
| | *CD276* | 0.2* | 0.3* |
| *Mostly downregulated genes* | *CD200* | -4.5 | -0.77 |
| | *TNFRSF14* | -2.9 | -6.7 |
| | *LGALS3* | -2.4 | -5.1 |
| | *BTN2A2* | -0.7 | -0.7 |
| | *CD48* | -2.3* | -2.5 |
| | *CTLA4* | -1.1* | -7.2 |
| | *VTCN1* | -1.3* | -5.0 |
| | *ADORA2A* | -1.4 | -1.7 |
| | *BTLA* | -1.4* | 0.0* |
| | *TIGIT* | -0.2* | 0.0* |
| | *BTNL3* | 0.0* | 0.0* |

All changes were statistically significant (p-values ranged between $p < 0.05$ to highly significant) unless marked by asterisk.

**Supplementary Table 2: Expression of co-stimulatory immune checkpoints genes in Rho $^{high}$ human melanoma cells.** RNAseq data of thirteen co-stimulatory genes of UACC62R relative to UACC62P and SK-Mel147 relative to SK-Mel-19

| | Gene | UACC62_R (Log2 fold change) | SK-Mel-147 (Log2 fold change) |
|---|---|---|---|
| *Mostly upregulated genes* | *CD86* | 7.7 | 0.0* |
| | *TNFSF18* | 5.2 | -2.3* |
| | *TNFSF4* | 1.3 | -1.8* |
| | *PVR* | 0.6 | 2.3 |
| | *NECTIN1* | 0.5 | 1.2 |
| | *BTN3A1* | 0.5 | 0.2* |
| | *BTN2A1* | 0.3 | 0.6 |
| | *HHLA2* | 0.6* | 0.0* |
| *Mostly downregulated genes* | *TNFRSF14* | -2.9 | -6.6 |
| | *ICOSLG* | -0.6 | -2.3 |
| | *CD40* | -2.3* | -7.5 |
| | BTNL9 | 0.0* | -4.3* |

All changes were statistically significant (p-values ranged between $p < 0.05$ to highly significant) unless marked by asterisk.

**Supplementary Figure S1**

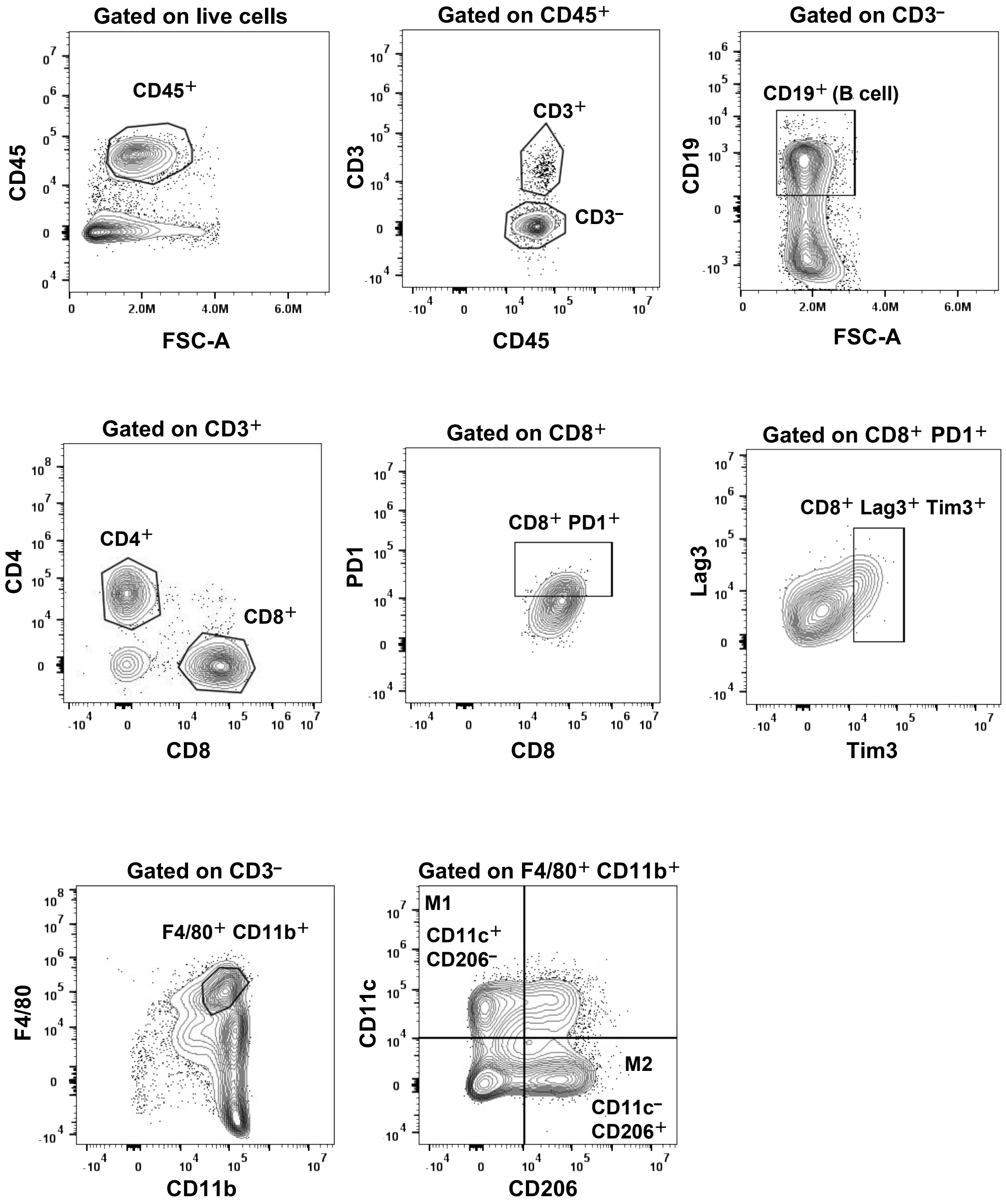

**Supplementary Figure S2**

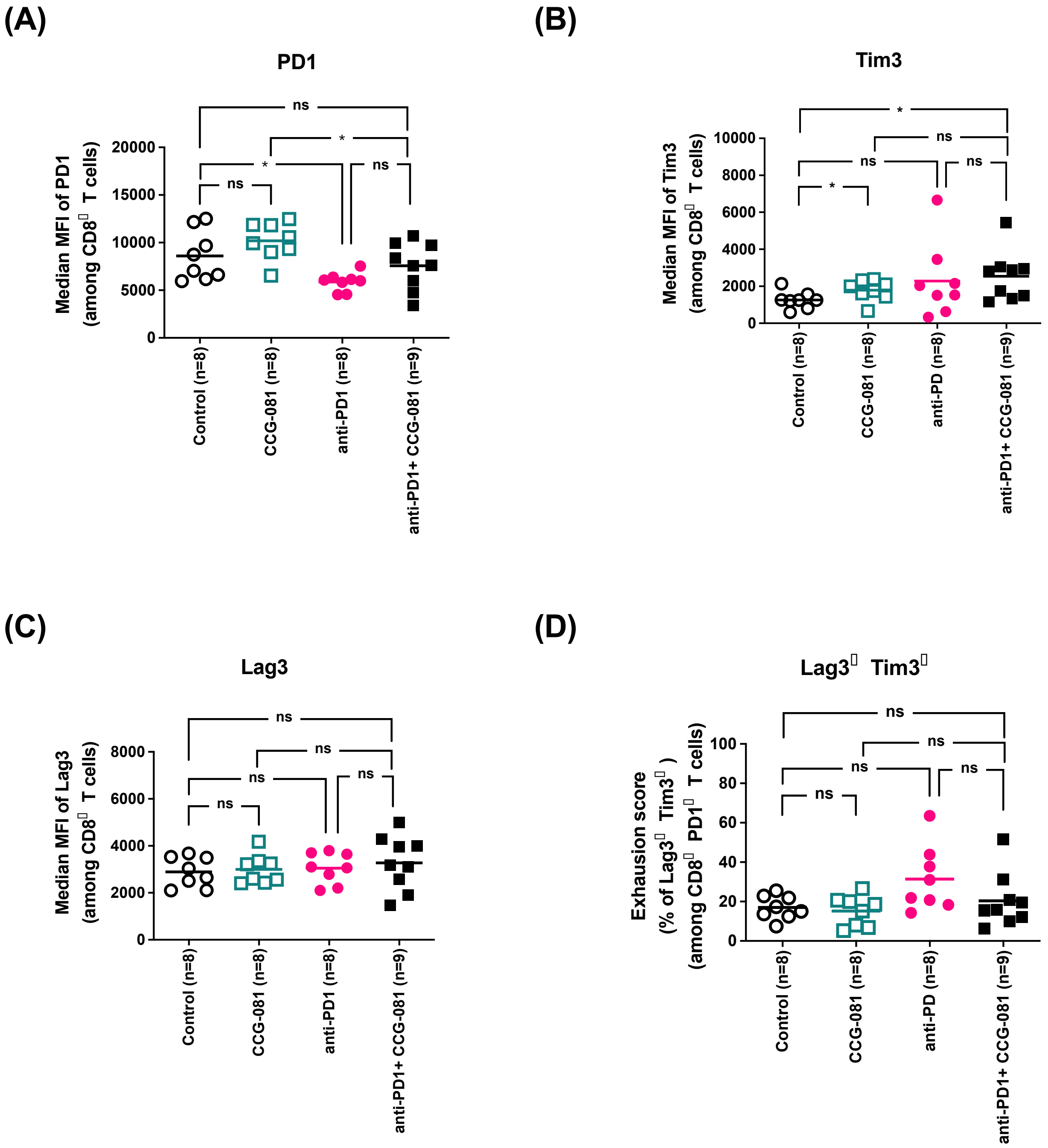
